# Breaking the next Cryo-EM resolution barrier – Atomic resolution determination of proteins!

**DOI:** 10.1101/2020.05.21.106740

**Authors:** Ka Man Yip, Niels Fischer, Elham Paknia, Ashwin Chari, Holger Stark

## Abstract

Single particle cryo-EM is a powerful method to solve the three-dimensional structures of biological macromolecules. The technological development of electron microscopes, detectors, automated procedures in combination with user friendly image processing software and ever-increasing computational power have made cryo-EM a successful and largely expanding technology over the last decade. At resolutions better than 4 Å, atomic model building starts becoming possible but the direct visualization of true atomic positions in protein structure determination requires significantly higher (< 1.5 Å) resolution, which so far could not be attained by cryo-EM. The direct visualization of atom positions is essential for understanding protein-catalyzed chemical reaction mechanisms and to study drug-binding and -interference with protein function. Here we report a 1.25 Å resolution structure of apoferritin obtained by cryo-EM with a newly developed electron microscope providing unprecedented structural details. Our apoferritin structure has almost twice the 3D information content of the current world record reconstruction (at 1.54 Å resolution ^1^). For the first time in cryo-EM we can visualize individual atoms in a protein, see density for hydrogen atoms and single atom chemical modifications. Beyond the nominal improvement in resolution we can also show a significant improvement in quality of the cryo-EM density map which is highly relevant for using cryo-EM in structure-based drug design.

## Introduction

Cryo-electron microscopy (Cryo-EM) has become very popular and successful in solving three-dimensional (3D) structures of macromolecular complexes. Improvements in electron microscopical hardware, image processing software and computing were the basis of what is known today as the resolution revolution in cryo-EM^2^. The recent years saw a stunning exponential growth in high-resolution structure determination of macromolecular complexes (https://www.ebi.ac.uk/pdbe/emdb/statistics_main.html) accompanied by a continuous shift in the attainable highest resolution of structures. Whereas the majority of the solved protein structures is still in the 3-4 Å resolution regime, a growing number of structures now reaches the 2-3 Å range, and very few examples are at a resolution better than 2 Å. The current record in the field is a 1.54 Å resolution structure (EMD-9865) of apoferritin ^1^. This structure was determined using a Jeol CryoARM 300 microscope equipped with a cold field emission gun electron source and an energy filter^3^. Obtaining increasingly higher resolution raises important questions about the ultimate resolution limit of single particle cryo-EM. How far can we push this technology and what are the limiting factors on the instrumental side? Is better hardware even required, or is it possible to overcome optical limitations by image processing software alone? Our goal was to address these questions by solving the structure of a biological sample at true atomic resolution and therefore directly visualizing all atoms in a protein, including the hydrogen atoms.

### New electron microscope hardware

We have used a prototype instrument equipped with additional electron-optical elements to increase the performance of the electron microscope. A monochromator ^4^ and a second-generation spherical aberration corrector ^5^ (CEOS GmbH) were built into a Thermofisher Titan Krios electron microscope equipped with a Falcon 3 direct electron detector. This hardware combination improved the optical properties by both a reduced energy spread of the electron beam, as a result of the monochromator, and a reduction of optical aberrations such as axial and off-axial coma, by the aberration corrector. As a reference, the microscope used here has an energy spread of about 0.1 eV, which is smaller than microscopes equipped with either Shottky field- (≈0.7 eV) or cold field-emission electron sources (≈0.4 eV), providing significantly increased temporal coherence and less dampening of high-resolution structural details in the images (Fig. 1). The additional BCOR spherical aberration corrector provides images that are free of axial and off-axial coma, which in the 1 Å resolution regime is the most limiting aberration. Additionally, the BCOR corrector can correct aberrations up to the fifth order and can also be used to minimize linear distortions in the images which result in non-uniform magnification of the images (Supplemental Figure 2f). Linear distortions are always present in electron microscopy and high-end microscopes typically suffer from linear distortions, which are in the range of 0.5-1%. Such values are negligible in attaining 3 Å resolution structures for relatively small specimens, but can become resolution limiting for larger macromolecular complexes and when approaching atomic resolution. Typical linear distortions in our microscope can be minimized to 0.1% and remain stable over longer microscope operation times. Images obtained from this new microscope thus neither require subsequent correction of linear distortions nor of coma by means of image processing as is usually the case for images obtained from standard electron microscopes. Nevertheless, as a consequence of the data collection over several months, we observed a small change over time in the overall magnification which we had to correct for (Supplemental Figure 2g). For projects where data can be collected within a few days, such magnification changes do not occur, but it is both essential and non-trivial to determine the exact magnification for reliable atomic model building and refinement.

**Fig.1.**
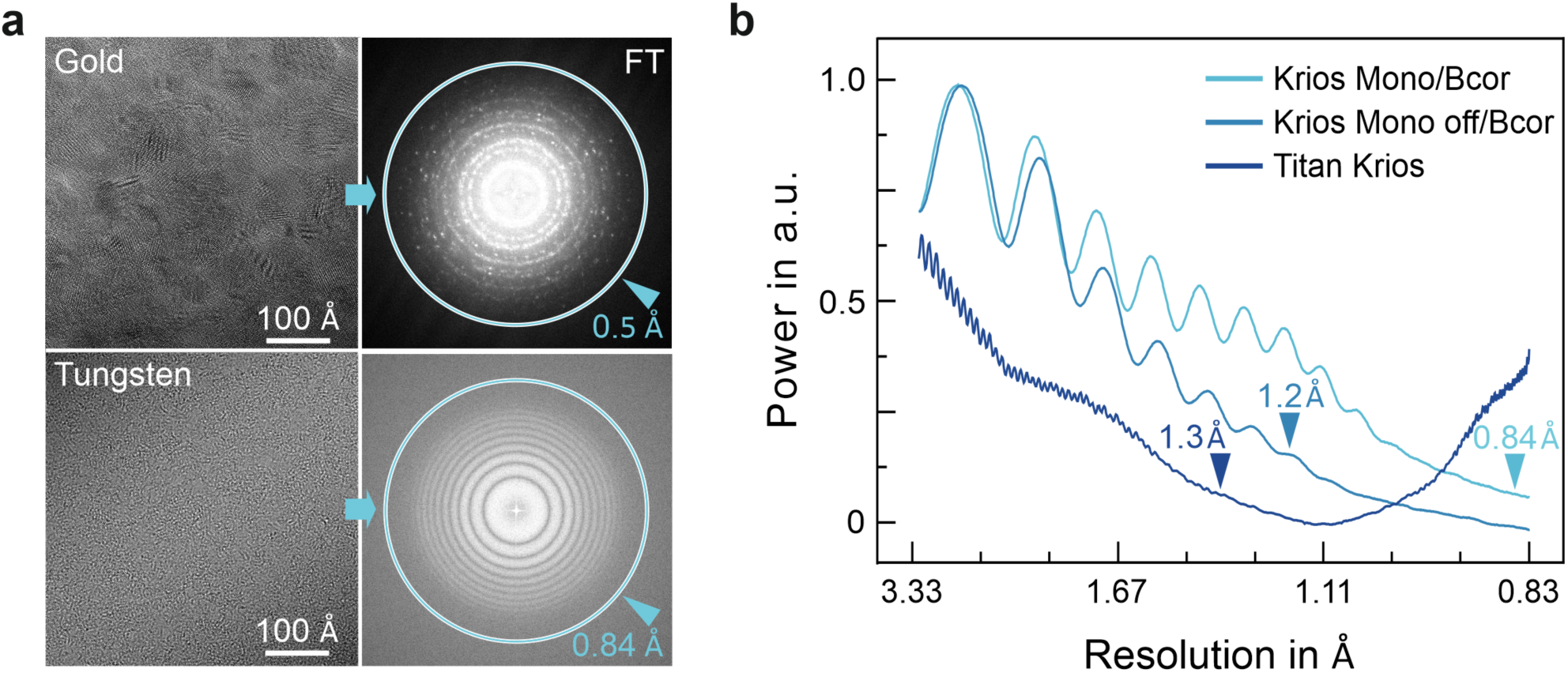
Optical performance of the Titan BCor/Mono electron microscope. a) High-resolution information transfer beyond 1 Å using a monochromator and Bcor corrector. Top: High-resolution image of a gold cross-grating specimen (Gold) showing reflections up to 0.5 Å in the Fourier transform (FT). Bottom: Image of a Tungsten specimen (Tungsten) and corresponding power spectrum (PS) showing information up to 0.84 Å according to equi-phase averaging as implemented in GCTF ^17^. The FT and PS were cropped differently for better visualization. b) Performance of the Krios Mono/Bcor vs. the system without monochromator (Mono off) and a standard Titan Krios. Power spectra from Tungsten obtained by equi-phase averaging. The maximum estimated resolution by Gctf is indicated (arrow-heads). See Methods for details.

Overall, the monochromated- and aberration corrected- electron microscope (Titan Mono-BCOR) showed an improved optical behavior reaching 0.5 Å resolution on a gold crystal (Fig. 1) in all spatial directions. This is a significant improvement in electron optical resolution of about 0.4 Å over a non-corrected- and non-monochromated-Titan Krios electron microscope. While the instrument clearly improves in optical resolution, how this translates into advances in the resolution attained for protein structures remains unclear. Multiple factors impede achieving higher resolution in structures of biological specimens, the quality of the sample itself being the most dominant ^6^. Biological molecules can be damaged during biochemical purification or the grid preparation process. Additionally, a multitude of dynamic conformational states, a characteristic of macromolecular machines, are imaged. This often restricts attaining high-resolution as image classification methods sometimes fail to classify different conformational states accurately. Essentially, when attempting to achieve higher resolution, the quality of biological specimens should be extraordinary.

### Resolution and Quality

Because of the beam sensitivity of biological objects, they are imaged at low electron dose and consequently the images suffer from a significant amount of noise. To fight this, an averaging procedure using several hundred thousand or sometimes millions of particle images is used to calculate one 3D structure by image processing. How many such particle images are needed to obtain a specific resolution can be described by an experimental B factor (as defined by Henderson and Rosenthal ^7^). The relationship between particle numbers and resolution becomes linear when plotting the logarithm of the particle number against the reciprocal square of the resolution and the experimental B factor is determined from the slope of this curve (Fig. 2). The experimental B factor represents a summary of all resolution limiting factors of a given electron microscope and describes the overall quality of the instrumental setup. Additionally, the B factor plots enable the extrapolation of how many particles will be required to reach even higher resolution (Fig. 2b), assuming that exclusively image noise is crucial and no other resolution-limiting factors exist. However, this assumption is crude, as additional hardware problems and aberrations are likely to occur when aiming for higher resolution.

**Fig. 2.**
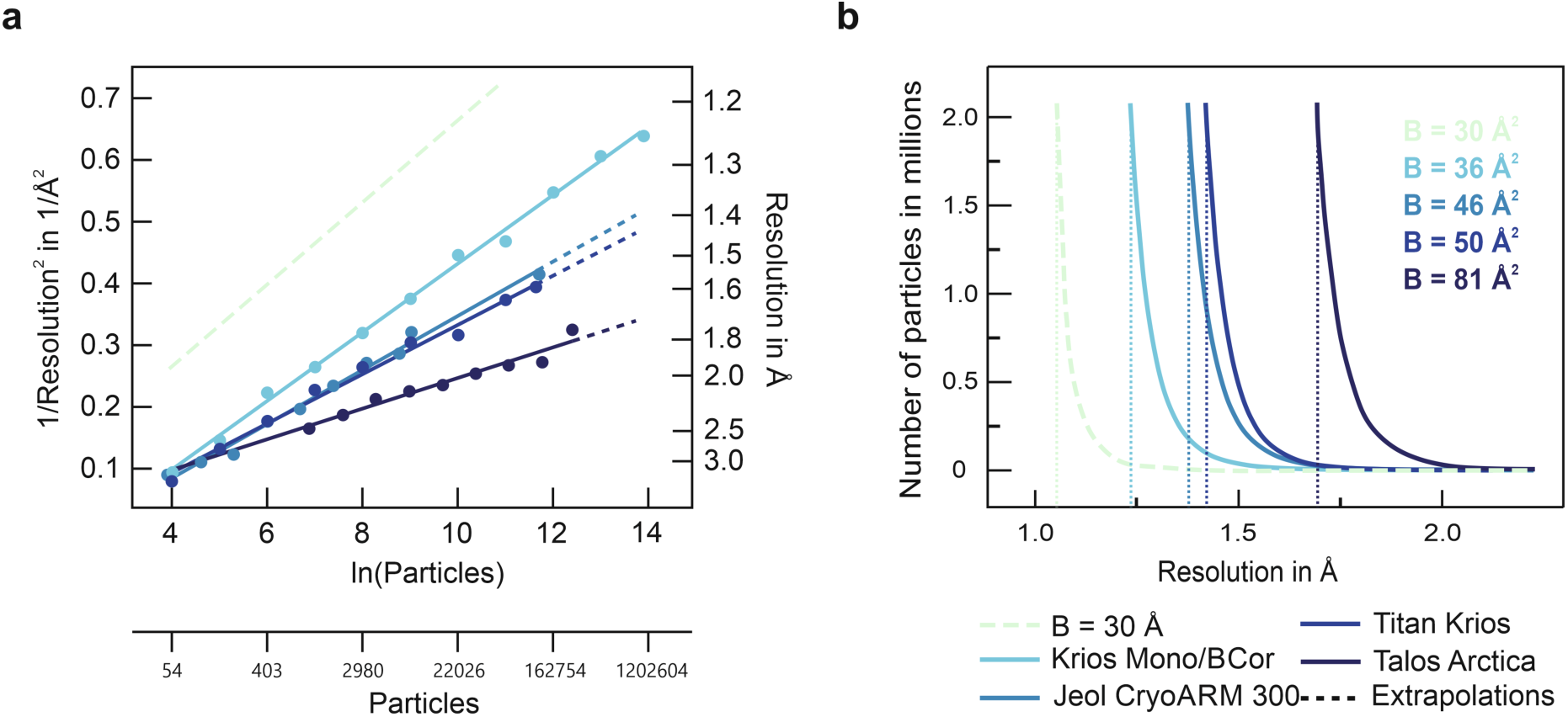
Performance comparison of electron microscopes. Performance comparison of cryo-EM systems. a) Performance as measured by the number of apoferritin particle images required to achieve a certain resolution cryo-EM structure. Remarkably, the Krios Mono/Bcor achieves superior performance without the need for *in silico* aberration-correction. B=30 Å^2^, plot for a theoretical B-Factor of 30 Å^2^: Krios Mono/Bcor, present data; Jeol CryoARM 300: EMPIAR-10248 & EMD-9865 ^1^; Titan Krios: re-computed from EMPIAR-10216 (see Methods); Talos Arctica: EMPIAR-10337 & EMDB-21024 ^23^. b) Experimental B-factors result in hard resolution-limits for a given cryo-EM imaging system. The explosive nature of required particle numbers at high resolution becomes more readily apparent by displaying B-factors in a linear plot. Note the similar performance of the various microscope systems at resolution up to 2 Å. The B factor of 36 Å^2^ obtained for the Krios Mono/Bcor indicates to become limiting for statistical particle number requirements only when attempting to attain structures slightly better than 1.25 Å. A microscope that could break the 1 Å resolution barrier with acceptable particle number statistics needs to have a B factor of 30 Å^2^ or better.

Summarizing what we currently know from existing state-of-the-art commercial EM hardware and their B factors, we can predict that several hundred billion apoferritin particle images would be required to reach 1 Å resolution (Fig. 2). With the current speed of data acquisition this would translate into several hundred years of data recording and a non-realistic amount of compute power and data storage capacities. These numbers clearly indicate that 1 Å resolution reconstructions are beyond any realistic expectations in cryo-EM using even the currently best commercially available microscope hardware.

In this manuscript, we try to answer several important questions such as: (i) What is the resolution limit that can be achieved by cryo-EM using existing technology? (ii) What are the particle numbers needed for the highest attainable resolution with our instrument described above? (iii) Is there any further technological improvement necessary and useful to push cryo-EM resolution boundaries? and (iv) What features are visible in cryo-EM maps at the highest obtained resolution using our advanced instrument?

### High-resolution structure of apoferritin

To answer these questions, it is important to consider which structural features are to be expected at a given resolution in EM. Structures of proteins in the 2-3.5 Å regime published so far show very comparable features as known from X-ray crystallography, even though the scattering of electrons and X-ray photons are substantially different physical events. However, this is expected to change in the very high-resolution regime. In X-ray crystallography, it is particularly challenging to see hydrogen atoms due to the limited photon scattering power of the single electron in a hydrogen atom 8. The situation in cryo-EM is different because electrons are scattered by the nuclear potential, which –in the case of H atoms-results in a significantly larger scattering cross section ^9^ compared to photons in X-ray crystallography ^10^. Whereas H-atoms become visible in X-ray maps typically only at resolutions close to 1 Å or better, they are supposed to become visible at somewhat lower resolution in cryo-EM.

To attain a high-resolution structure of apoferritin, we recorded images using gold grids (1.2µm and 1.2µm holes), a pixel size corresponding to 0.492 Å and a total dose of about 40 electrons per Å^2^, on a Falcon 3 detector in electron counting mode (Suppl Fig. 1). The B factor plot calculated based on the initial data indicated that more than 5 million apoferritin particle images would be required to attain a resolution of 1.3 Å, i.e. a resolution where hydrogen atoms might be visualized due to the differences in H-atom cross section between photons and electrons discussed above. Optimization of grid preparation and imaging conditions allowed us to cross the 1.5 Å resolution barrier with only 17.800 particle images (B factor of 36 Å^2^). Using a total number of 1.000.000 particle images, we finally obtained a map at 1.25 Å resolution. This map – in contrast to X-ray maps at this resolution-shows well-defined additional densities that agree with the positions of hydrogens on almost all atoms (Fig. 3 and Supplemental Figures 3 and 4), when using low thresholds for map visualization. At higher thresholds, almost completely separated densities for individual C, O and N atoms are visible in most areas of the map (Fig. 3). Furthermore, the resolution of the map is sufficient to observe a sulfur chemical modification in apoferritin (Fig. 4), which to the best of our knowledge has not been visualized as yet.

**Fig. 3.**
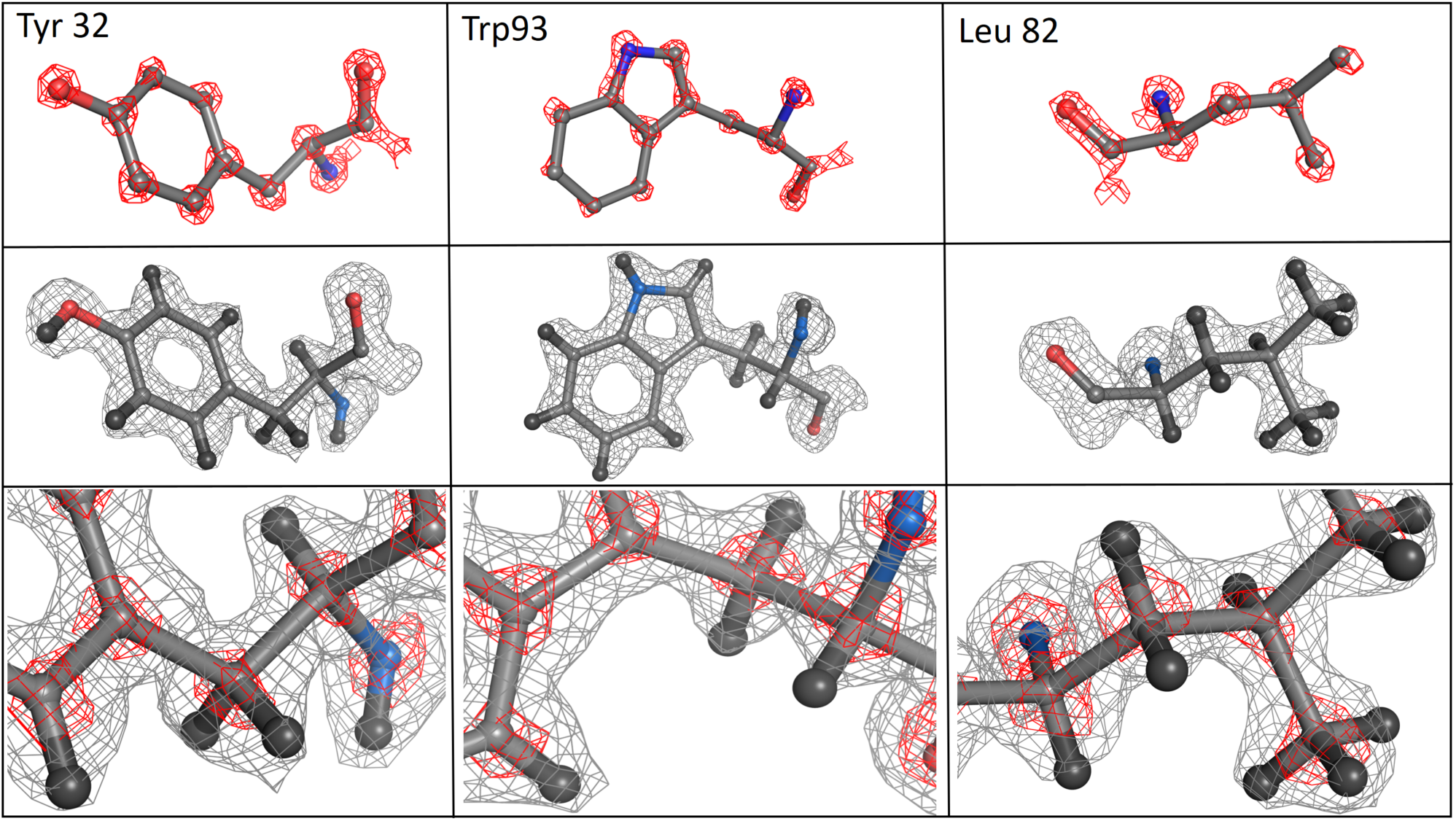
True atomic resolution structure of apoferritin – some structural details. True atomic resolution: Visualization of individual atoms and hydrogens at 1.25 Å resolution Three apoferritin residues are shown at high (red mesh) and low (grey mesh) density thresholds. The true atomic resolution of our map is shown in the first row by the clear separation of individual C, N and O atoms at high thresholds. The second row shows density for hydrogen atoms in all parts of the individual amino acid side chains. A ball and stick representation for the hydrogen atoms (in black) is included in the atomic model. Similar visibility of density for hydrogen atoms requires about 1 Å or better resolution in X-ray crystallographic structures (Supplemental Figure 3). The third row shows close-up views of the three amino acids using both density thresholds simultaneously.

**Fig.4.**
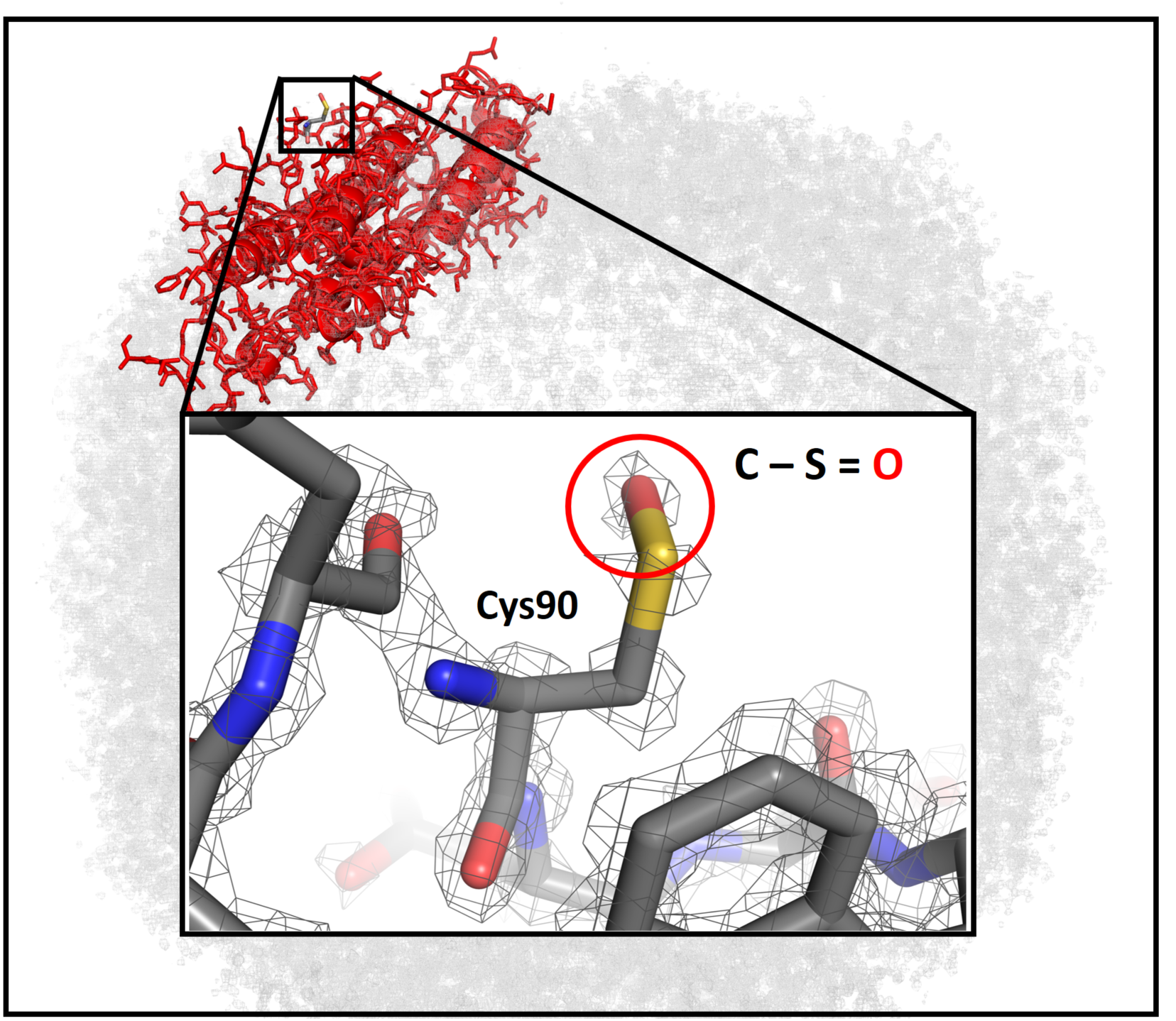
Visualization of a single atom chemical modification. Cysteine 90 of human apoferritin is located at the surface of the macromolecular complex. The solvent-exposed regions are usually determined at lower local-resolution in cryo-EM but in our high-resolution structure it is still sufficient to visualize a single atom oxygen modification (marked by a square on the entire molecule and by the red circle in the close-up).

### Resolution vs. Map Quality

How do the various cryo-EM structures for apoferritin compare both in terms of required particle statistics and map quality? In cryo-EM, the resolution of the map is estimated by the correlation of two independently calculated structures in various resolution shells in Fourier space (Fourier shell correlation ^11^). The FSC provides a single number for the obtained resolution, but does not provide a direct measure of the quality of a 3D structure. It is also noteworthy that the resolution estimated by the FSC is not entirely independent from data processing procedures and indeed some optical aberrations such as coma can potentially lead to resolution overestimation^12^ and generation of map artefacts. Therefore, an independent means of map quality comparison, emphasizing quality in addition to the FSC-based resolution estimation is desirable. In principle, the degree by which the map can be interpreted in atomic modelling, particularly with respect to the bound solvent and subsequent refinement of the model, represents such a measure. Atomic model refinement at intermediate resolutions strongly rely on prior knowledge of (protein-) chemistry such as bond-lengths, - geometry and additional constraints such as the planarity of aromatic systems. At atomic resolution, the density of information gleaned from the structure should allow for atomic modelling independent of constraints and enable the experimental observation of slight deviations from standard chemistry. Such deviations in fact frequently arise and are essential aspects in understanding how proteins catalyze seemingly impossible reactions. Sub Å resolution crystallography reveals such distortions that enhance enzyme reactivity of enzymatic intermediates ^13,14^. In addition, such detailed, experimental and unbiased views of protein architecture can provide invaluable insights into binding pockets for the design of specific drugs. In the case of cryo-EM density maps, features deviating from perfect geometry can either reflect such true changes or they might be caused by image processing errors and/or optical aberrations. While a certain deviation from perfect geometry is allowed, a good quality high-resolution cryo-EM map should allow the modelling of proteins with no overall systematic deviations from normal chemical structure and allow the refinement of atomic models against the experimental map in an unconstrained fashion. Other quality estimators are the abundance of ordered solvent, which can be modelled, and the coordination distances of ordered solvent molecules from the protein. Both these traits of ordered solvent can be expected to increase with the quality and resolution of the determined experimental structures.

We therefore compared a 1.55 Å resolution structure from a subset of the data acquired in our instrument with the highest resolution apoferritin structure at 1.54 Å, which should allow for the assessment of quality, independent of resolution differences (Suppl Figure 5). Notably, to achieve this resolution in our microscope, only about 17.800 particle images were required that could be recorded in a single day. This is roughly seven times less data compared to the particles needed to achieve 1.54 Å using the Jeol microscope ^1^. Even though the nominal resolution is comparable, we were able to reliably model 1.750 more water molecules in our map applying the same procedures (Supplemental Figure. 5). This significant difference in map quality can be attributed to the improved optical quality of the Titan Mono-BCOR microscope. Whereas no beam-tilt induced coma is present in our data, there is substantial amount of beam tilt present in the data recorded in the Jeol CryoARM microscope due to some instabilities of the cold field emitter after flushing. Latest developments in the Relion software ^15,16^ allow the *a posteriori* computational correction of beam tilt induced coma, the accuracy of these corrections however are unknown.

### Can we expect further improvements by new EM hardware?

For perfect samples the resolution and the quality of the 3D density map are dependent on the electron microscopic hardware and we show here that atomic resolution is attainable. For the vast majority of macromolecular complexes, resolution improvements will additionally depend very strongly on improvement in sample quality as well as on image classification tools that can handle continuous conformational motion in the data. Even though it is possible to obtain 1.5 Å resolution with 17.800 particles only, it is noteworthy that in case of a non-symmetric and more dynamic complex the required particle numbers can be several orders of magnitudes higher. For asymmetric particles, adopting 10 conformational states that need to be computationally sorted, would already require the acquisition of an unrealistic number of >5 million particles using the Titan Mono-BCOR microscope. Any improvement in image recording speed by faster cameras and optimized data acquisition schemes will contribute substantially to attain such goals in the future. Realistically, cryo-EM structure determination at resolutions that allows the visualization of individual atoms and hydrogens is not likely to be possible in high-throughput in the near future. However, if resolutions in the 1.5-2 Å range are sufficient, a microscope as presented here can be powerful enough to determine several structures a day for a biochemically well-behaved and symmetric macromolecular complex. At slightly lower resolution aims, the microscope itself becomes an increasingly non-decisive factor and in the 3-3.5 Å resolution range, all the currently commercially available microscopes with a direct electron detector can be used because the required particle numbers do not depend primarily on the experimental B factor of the instrument (Fig. 2a). Each electron microscope does have strict limits in its high-resolution capabilities because of the exponential nature of the required particle numbers once the optical limits of the systems are reached (Fig. 2b). Our current microscope has an experimental B factor of 36 Å^2^ which makes it almost impossible to obtain a resolution better than 1.2 Å. However, next generation detectors are likely to improve speed and quality of image detection. It remains to be seen whether such new hardware could help to push the B factor to a value of <30 Å^2^ which would be required to break the 1 Å resolution barrier using a manageable amount of data (roughly 2 million apoferritin particles).

Currently available electron microscopes have very hard limits in their maximum resolution capabilities that can hardly be overcome by image numbers (Fig. 2). Improving electron microscope hardware is therefore essential for a more quantitative level of protein structure interpretation in cryo-EM at very high resolution. This work shows a significant step forward into this direction providing protein structures at an unprecedented resolution allowing atomic details to be investigated. To reach true atomic resolution on a more regular basis and higher throughput one will require further developments in cryo-EM technologies, which is realistically to be expected in a field that develops so quickly. The next generation electron microscopes and detectors have the potential to make a significant contribution to our understanding of catalytic mechanisms of proteins and to the development of new drugs based on high resolution and high quality 3D structures of proteins.

## Material and Methods

### Apoferritin expression and purification

A cDNA encoding for the K86Q variant of the human ferritin heavy chain with codons optimized for E.coli expression was obtained from Geneart (Regensburg) and subcloned as a NdeI-XhoI fragment into the pRSET-A expression plasmid. The untagged protein was obtained by transformation of this plasmid into BL21(DE3). Expression was performed in Terrific Broth medium containing Ampicillin (100 µg/ml) and Glucose (1 % w/v), cells were induced at OD_600_ = 0.8 with 1 mM IPTG for 3.5 hours at 37 °C and the harvested cells were then stored at −20 °C. To purify the protein, the cell pellet was resuspendend in 15 mM Tris-HCl (pH 7.4), 150 mM NaCl, 10 mM DTT, 10 mM Benzamidine-HCl, 10 mM EDTA, 1 mM PMSF, 2.5% (w/v) sucrose, lysed by the addition of 1 % Triton X-100 and passage through a high-pressure disruptor (Emulsiflex). A cleared lysate was prepared by centrifugation at 30.000 x g, followed by the precipitation of nucleic acids by the addition of 3 % (w/v) streptomycin sulfate and centrifugation at 20.000 x g. The supernatant was incubated at 75 °C for 20 minutes to precipitate E. coli proteins, followed by centrifugation at 20.000 x g for 30 minutes. The resulting supernatant was then subjected to ammonium sulfate fractionation by the addition of powder to a saturation of 60 % (w/v). The resulting pellet was resuspended in 15 mM Tris-HCl (pH 7.4), 150 mM NaCl, 10 mM DTT and 0.1 % (w/v) LMNG and subjected to density gradient centrifugation on 10 - 40 % (w/v) sucrose gradients in a Surespin CA 630-36 rotor at 25.000 rpm, 18 °C for 19 hours. Fractions containing Ferritin were identified by SDS-PAGE, pooled and subjected to dialysis against 50 mM MES, 150 mM NaCl, 0.5% thioglycolic acid (pH 6.5) for 24 hours, followed by dialysis against 15 mM Tris-HCl (pH 7.4), 150 mM NaCl and 2.5% (w/v) sucrose for 24h at room temperature. The resulting apo-ferritin was then precipitated by the addition of ammonium sulfate to 60 % saturation and the pellet resuspended with 15 mM Tris-HCl (pH 7.4), 150 mM NaCl, 1 mM EDTA and 0.01% (w/v) LMNG. The sample was then subjected to density gradient centrifugation on 10 - 40 % (w/v) sucrose gradients in a SW40Ti rotor at 30.000 rpm, 18 °C for 16 hours. Fractions containing Ferritin were identified by SDS-PAGE, pooled and buffer exchanged to 1 x PBS by concentration in an Amicon spin concentrator and frozen in liquid nitrogen and stored in small aliquots at a concentration of 10 mg/ml.

### Cryo-EM grid preparation

For alignment purposes, UltrAuFoil R1.2/1.3 and R1.5/1.6 300-mesh grids (Quantifoil, Jena) were pre-floated with custom-made continuous carbon foil covering ∼25% of the grid area. Grids were glow discharged for 10 seconds with a glow discharger built in-house under low vacuum shortly prior to sample application. 4 μL of purified apoferritin (∼3.5 mg mL^-1^) was applied to glow-discharged grids, which were blotted and plunge-frozen with a Vitrobot Mark IV (Thermo Fisher Scientific, Eindhoven) operated at 4°C and 100 % humidity; the blotting time was set to be 6.5 seconds and 7.5 seconds for UltrAuFoil R1.2/1.3 and R1.5/1.6 grids, respectively.

### Cryo-EM data acquisition and image processing

All cryo-EM data were collected in nanoProbe mode on a Titan Krios electron microscope operating at 300 kV equipped with a monochromator (Thermo Fisher Scientific, Eindhoven) and an aplanatic image corrector “B-Cor” (CEOS, Heidelberg). The monochromator was tuned to operate at 3 kV potential and excitation (0.8) to obtain an energy spread of the source of about 0.1-0.15 eV. The B-Cor was set-up to correct for off-axial aberrations, and beam-image shift induced electron optical aberrations coma and for linear distortions. For each cryo-EM data set, the Bcor was tuned with the signals from the amorphous carbon i) to correct the phase errors introduced by on-axis electron optical aberrations to less than 45 degrees at scattering angles ≥ 17 mrad, which is equivalent to ≤1.16 Å resolution, and ii) to reduce linear distortions to ≤0.2%.

A total of 10.398 movies with 40 fractions each were collected. The exposure time was over 18.54 seconds with a dose of ∼1.25 e-Å2 per fraction in electron counting mode using a Falcon III direct electron detector (Thermo Fisher Scientific, Eindhoven) at a nominal magnification of 120.000x (∼0.492 Å pixel-1) and 0.3 μm to 1 μm underfocus resulting in a total dose of ∼50 e-Å2. An adapted version of the software EPU (Thermo Fisher Scientific, Eindhoven) was used to acquire data by beam-image shift over up to 3 × 3 holes; for R1.2/1.3 grids three movies were acquired per hole, for R1.5/1.6 grids five movies per hole. The data was acquired over 40 acquisition sessions resulting in 40 data sets, which were each pre-processed independently.

All image processing was performed using RELION 3.1^16^, if not indicated otherwise. Global motion correction and dose-weighting were performed with a B-factor of 150 and the dose-weighted micrographs were used for CTF estimation by GCTF 1.06^17^. Unless otherwise specified, all subsequent image processing steps were per-formed in RELION 3.1. Particles were picked with reference to 2D templates and extracted with 2 × 2 binning (0.984 Å pixel-1, 180×180 pixel box). Two rounds of reference-free 2D classifications were performed to remove bad particles. The remaining good particles were re-centered, and re-extracted in full size in a 480×480 pixel box (0.492 Å pixel-1) and subjected to 3D refinement; all 3D procedures were performed imposing octahedral symmetry. The structure of mouse apoferritin (EMDB-9599, REF) was low-pass filtered to 20 Å pixel 1 and used as a reference for initial 3D refinements. A first round of CTF refinement was performed on the refined particles to correct for per-micrograph defocus and per-micrograph astigmatism. Then, particle motion and the corresponding per-frame relative B factors were estimated by Bayesian polishing and the polished particles were subjected to a second round of 3D refinement. Another round of CTF refinement was performed to correct for per-particle defocus and per-particle astigmatism. Afterwards, the particles were again subjected to Bayesian polishing and a third round of 3D refinement. The procedure was repeated for all 31 datasets independently.

Particles from individual data sets were defined as a unique optics group each and all 1.466.774 particles were combined for all further image processing. CTF refinements were performed to correct first for magnification anisotropy, then for 4th order aberrations (to refine the spherical aberration constants which varied slightly due to the B-Cor tuning) and, finally, for per-particle defocus, per-particle astigmatism, followed by another 3D refinement. Ewald sphere correction (REF) was performed on the two resulting half-maps yielding a final reconstruction with a resolution of 1.272 256 Å for a pixel size of 0.492 Å pixel-1, as calibrated by atomic-model refinement (see below). Subsequently, particle images were re-polished based on the new 3D structure and windowed and refined in a 600×600 pixel box to account for the high-resolution information-spread introduced by defocus. The refined particles were reduced to 1.090.676 particle images, excluding particles with a defocus >9000 Å and a maximum-value-probability distribution <0.04. From the reduced data set, half-maps were reconstructed using Ewald sphere correction ^18^ resulting in a 3D reconstruction at a final resolution of 1.25 Å (Supplementary Figure 2b).

### B factor plots

For the B-factor plot (Figure 2a), the total set of 1.090.676 particle images from the final refinement was randomly resampled into smaller subsets, half-maps were reconstructed for each subset using Ewald sphere correction and FSCs were computed by cropping the half-maps to 480×480 pixel boxes. For the standard Titan Krios, we re-evaluated data from EMPIAR-10216 mainly as described by the authors Radostin Danev and colleagues (https://twitter.com/radodanev/status/1022265459666694144?lang=en) but with modifications to at least partially account for off-axial aberrations by splitting the micrographs into 9 subsets^16^. This procedure slightly improved the final resolution from the stated 1.62 Å to 1.59 Å.

### Atomic model building and refinement

Only unsharpened maps were used for the refinement of models, as the utilization of sharpened maps resulted in severe distortions of model geometry. An initial model was generated using the deposited K69Q human ferritin heavy chain model (2CEI). The 24 mer for model refinement was assembled using MOLREP ^19^ anew into every experimental EM map using the 2CEI monomer stripped of all solvent. After rigid body refinement and positional refinement in Refmac5 ^20^ essentially as described ^21^, the model was adjusted to the density manually in Coot ^22^. After an additional round of positional refinement in Refmac5, solvent molecules were added interactively in Coot, followed by an additional round of refinement in Refmac5.

## Acknowledgements

We thank Jan Erik Schliep for his contribution in the early phase of the project and Martin Link and Sodkhuu Dalaikhuu for their technical contribution in the initial setup of the microscope. This work was supported by the Max Planck Society and the Deutsche Forschungsgemeinschaft (grants SFB 860 to H. S.).

## Author contributions

HS, KMY and NF set up the Krios Mono/Bcor microscope. EP and AC developed the purification strategy and EP purified the complex. KMY prepared grids and collected EM data. KMY and NF did the cryo-EM image processing analysis. AC built and refined the atomic models. The manuscript was written by HS with input from all authors. HS initiated and orchestrated the project.

## Data availability

The atomic models have been deposited in the Protein Data Bank (PDB) with the following accession codes: 1.55 Å structure XXXX and 1.25 Å structure XXXX. The cryo-EM maps have been deposited in the Electron Microscopy Data Bank as follows: 1.55 Åmaps (EMD-XXXX) and 1.25 Å maps (EMD-XXXX)

## Supplementary Information

**Suppl. Fig. 1.**
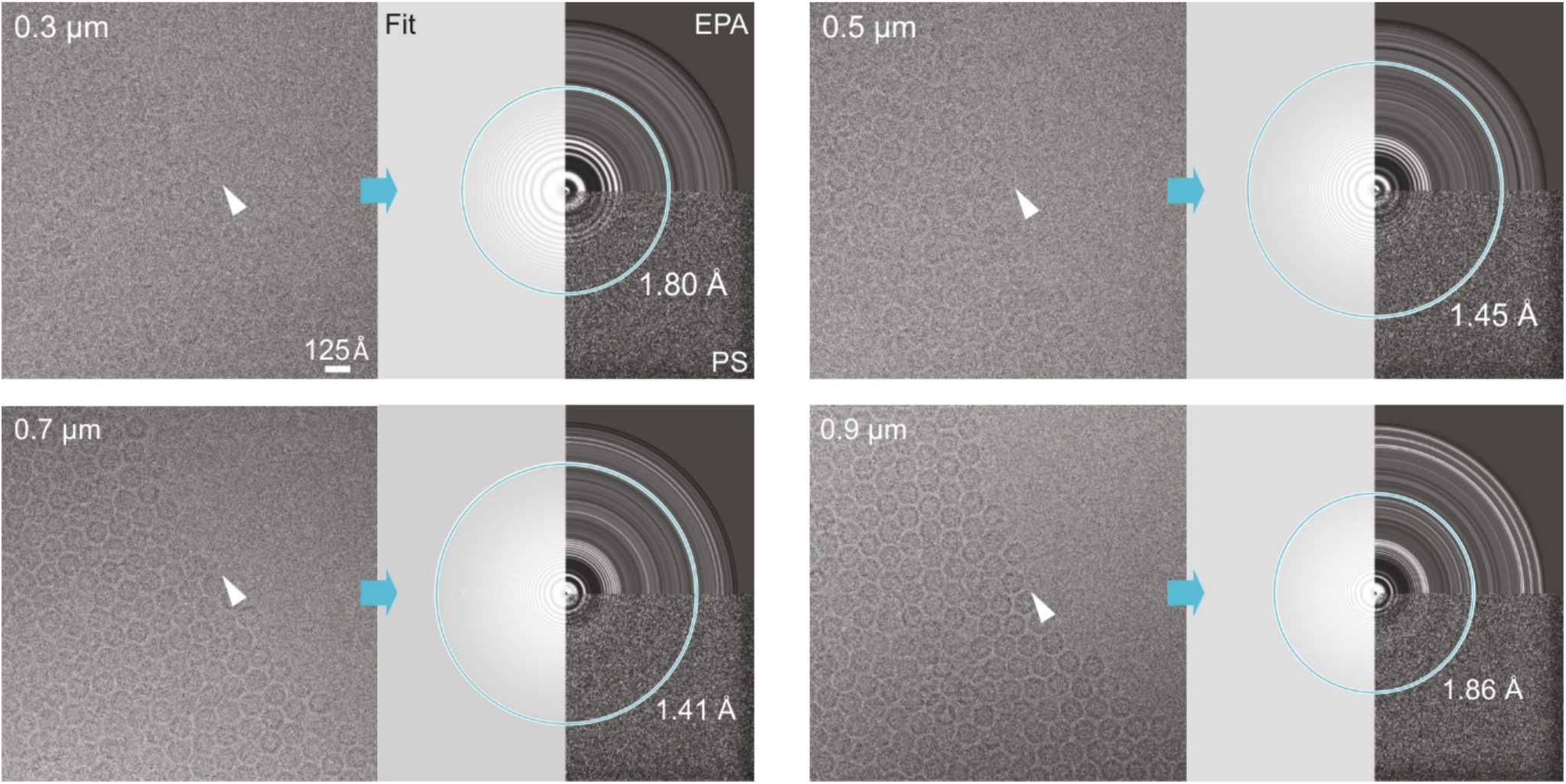
Low-dose cryo-EM micrographs of human apoferritin. Exemplary micrographs (left) acquired with a total dose of ∼40e-Å^2^ are shown with their power spectra (PS), equi-phase average (EPA) and the fit of the contrast transfer function (Fit). Numbers indicate the respective defocus in µm and the maximum resolution (in Å) in the power spectra as estimated by Gctf ^17^. White arrow heads denote the transition from areas with particles in dense packing to areas devoid of particles indicating a very thin layer of vitrified ice, as required for high-resolution imaging.

**Suppl. Fig. 2.**
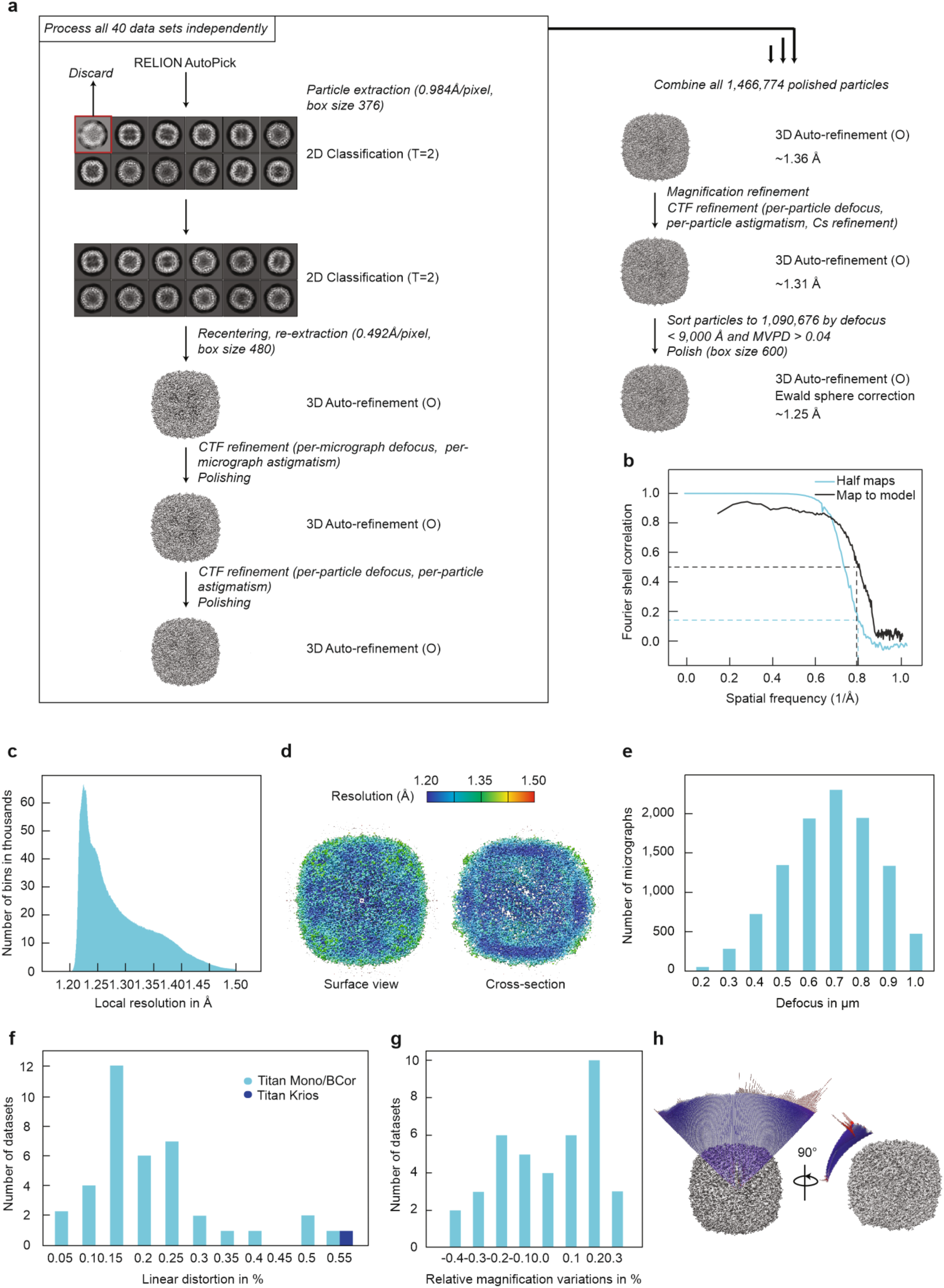
Cryo-EM structure determination. a) Image processing pipeline. See Methods for details. b) Fourier-shell-correlation plots for independently refined half-maps (Half-maps) and full map vs. model (Map to model). c) Histogram of local resolution for the final 1.25 Å map obtained with relion postprocess using a small soft spherical mask. d) Final map colored by local resolution. e) Defocus distribution for the total set of 10,398 micrographs. f) Linear distortions as estimated by relion_ctf_refinement for the present Krios Mono/Bcor data and data from a standard Titan Krios (EMPIAR-10216). g) Relative magnification variation in the present data as determined by relion_ctf_refinement. h) Angular distribution for the final map.

**Suppl.Figure 3:**
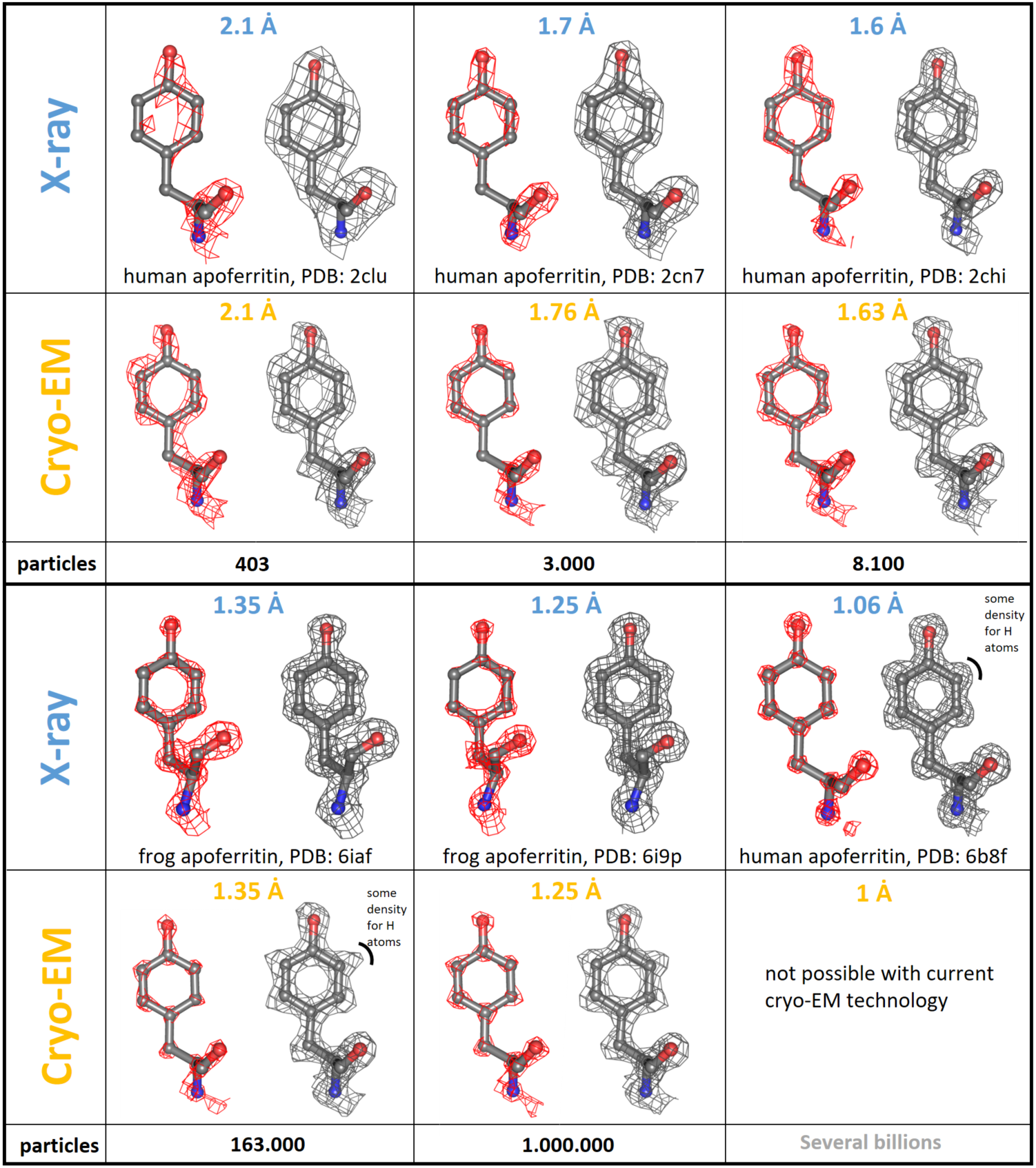
Comparing high-resolution features obtained by X-ray crystallography and cryo-EM. For the comparison, we selected the same tyrosine residue (Tyr 32 in human apoferritin, Tyr 34 in Frog) from our structures obtained at different resolutions with published crystallographic data of apoferritin at the indicated resolution. The same residue is shown at high (red) and low (grey) threshold to better judge the structural details and the map quality. Density that can be attributed to hydrogen atoms can barely be seen in any of the X-ray structures. Even at 1.06 Å resolution only weak density for hydrogens can be detected while the cryo-EM reconstruction already reveals some density for hydrogens at 1.35 Å resolution. At high thresholds, the separation into clearly distinct atoms can only be seen in the 1.06 Å resolution X-ray map but not in lower resolution X-ray data. In case of our cryo-EM reconstructions we can see individual atoms starting at 1.35 Å resolution or better. Cryo-EM structures at 1 Å resolution are currently not possible because they would require unrealistically high particle number statistics even with the Krios Mono/Bcor microscope (see Fig. 2).

**Suppl. Fig. 4:**
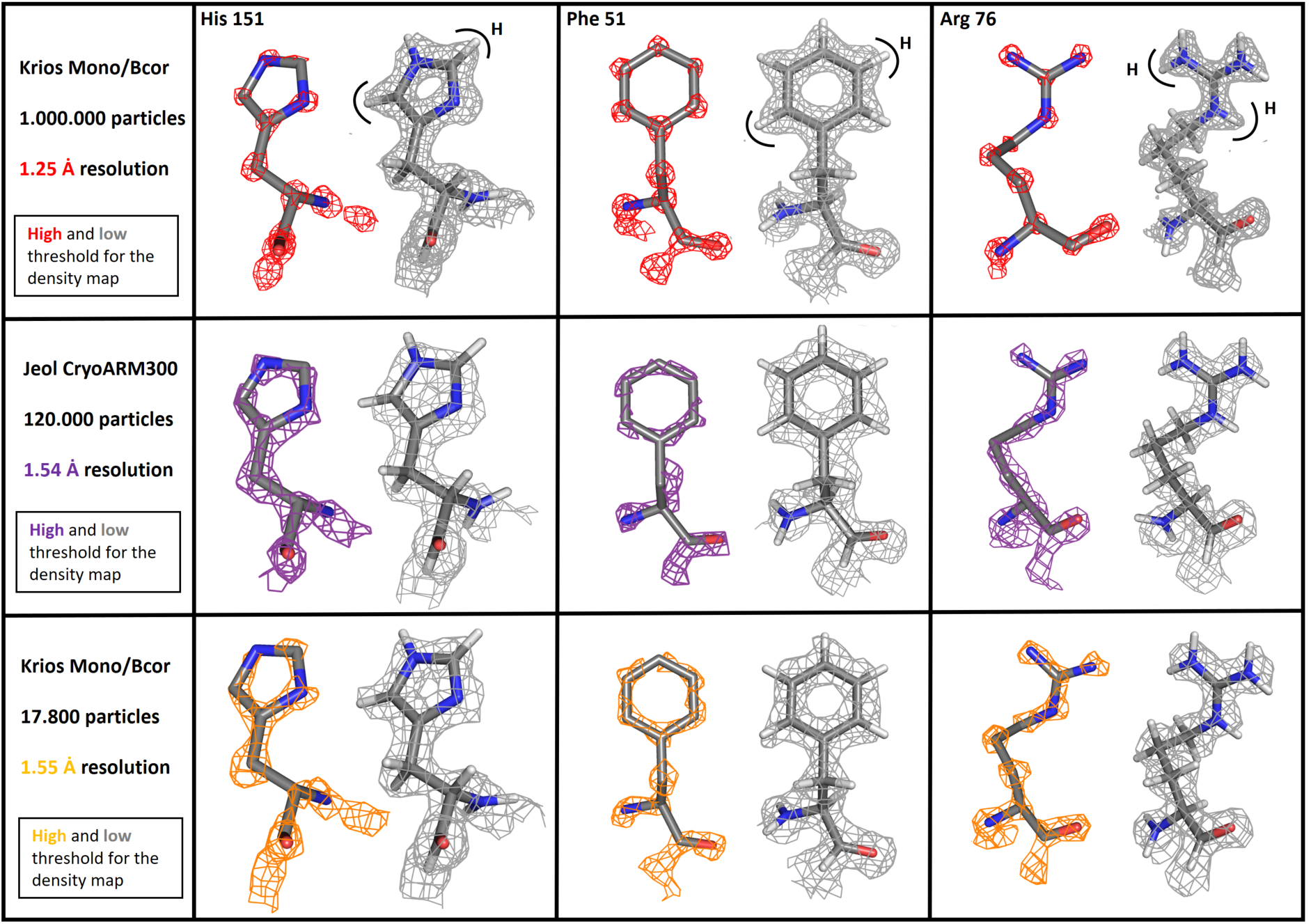
Structural features of our cryo-EM maps at 1.55/1.25 Ȧ resolution compared to the thus far reported highest-resolution map at 1.54 Ȧ resolution (EMDB-9865). Three apoferritin residues (His 151, Phe 51, Arg 76) are shown at two different density thresholds in three cryo-EM maps. Row one depicts our present high-resolution map at 1.25 Ȧ resolution and row three shows a structure at 1.55 Ȧ resolution obtained from a smaller subset of the same data. Only 17.800 particles were necessary for this reconstruction to obtain Ȧ resolution which is a 6.7x lower particle statistics compared to the 1.54 Ȧ Jeol CryoARM300 map (second row). The low-threshold density meshes are always shown in grey and H atoms (white sticks) are included in the corresponding atomic models. Only in the Krios Mono/Bcor structure at 1.25 Ȧ resolution density becomes visible to accommodate all hydrogen atoms. At higher thresholds the two structures at 1.54 Ȧ and 1.55 Ȧ resolution nicely maintain the shapes of the sidechains but only in the structure at 1.25 Ȧ resolution individual atoms become clearly separated from each other indicating true atomic resolution.

**Suppl Fig. 5.**
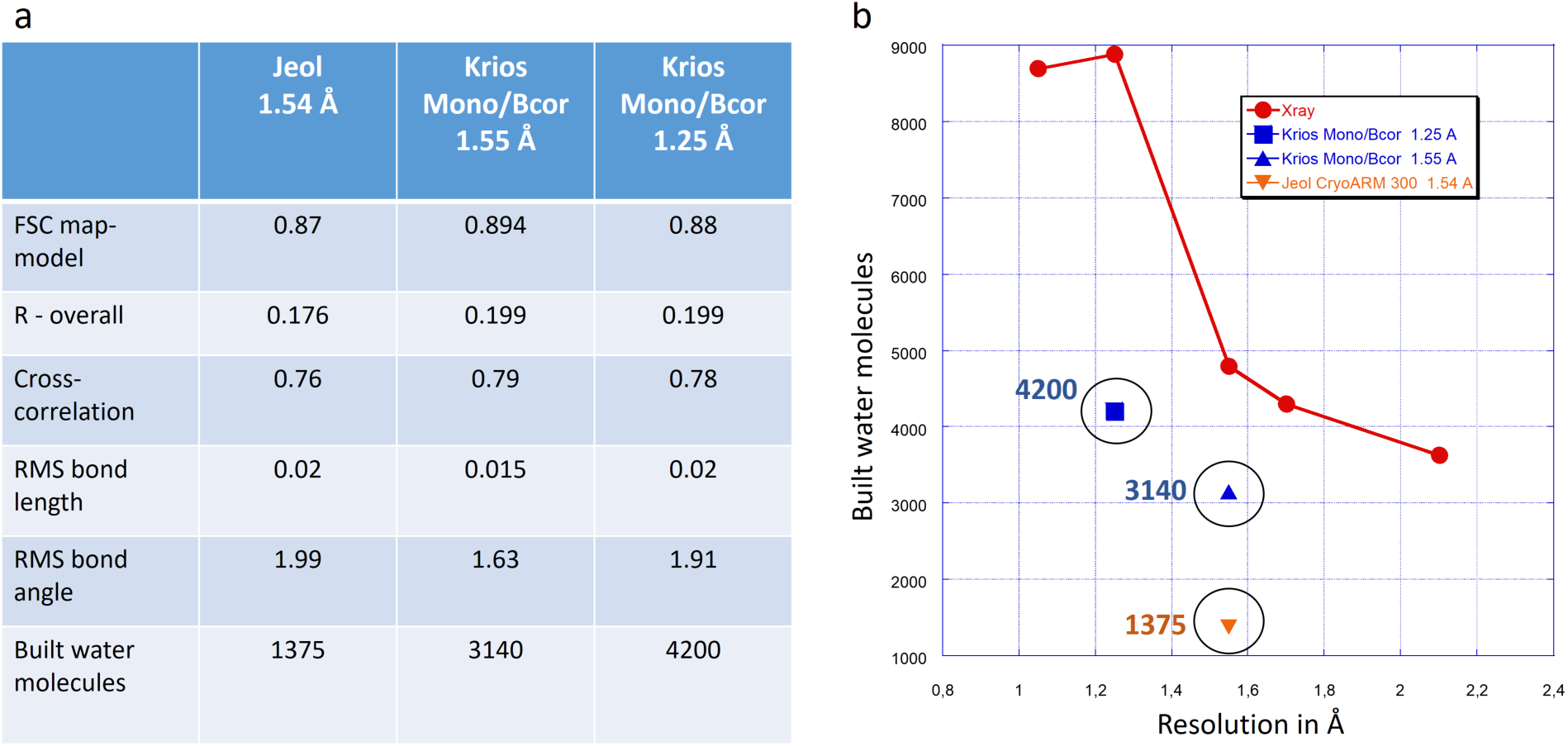
Relevant map-model building parameters and solvent molecules. a) As expected for such high-resolution structures, the calculated model validation parameters are in a very good range for all three cryo-EM map/models. The results from the two structures at 1.55/1.54 Å resolution can best be compared because they were determined at the same resolution. When comparing those two structures, all validation parameters are in favor of the Krios Mono/Bcor structure except the R value. The reason for that could be that 1765 more water molecules have been identified compared to the Jeol structure, which makes a detailed R value comparison difficult even though the two structures have been determined at the same resolution. b) The number of water molecules that can be localized in a structure rises with resolution and quality of a map, which is true for X-ray and for cryo-EM. The plot shows the number of water molecules found in the various X-ray structures (red dots) in relation to the three cryo-EM maps (blue and orange). Substantially more water molecules were identified for both Krio Mono/Bcor reconstructions compared to the Jeol map. It is noteworthy that the number of waters that were located in the X-ray structures is still higher than for the cryo-EM maps for reasons that are currently unknown.

